# Consensus Clustering for Robust Bioinformatics Analysis

**DOI:** 10.1101/2024.03.21.586064

**Authors:** Behnam Yousefi, Benno Schwikowski

## Abstract

Clustering plays an important role in a multitude of bioinformatics applications, including protein function prediction, population genetics, and gene expression analysis. The results of most clustering algorithms are sensitive to variations of the input data, the clustering algorithm and its parameters, and individual datasets. Consensus clustering (CC) is an extension to clustering algorithms that aims to construct a robust result from those clustering features that are invariant under the above sources of variation. As part of CC, stability scores can provide an idea of the degree of reliability of the resulting clustering. This review structures the CC approaches in the literature into three principal types, introduces and illustrates the concept of stability scores, and illustrates the use of CC in applications to simulated and real-world gene expression datasets. Open-source R implementations for each of these CC algorithms are available in the GitHub repository: https://github.com/behnam-yousefi/ConsensusClustering

## 1. Introduction

Clustering, or cluster analysis, is a widely used technique in bioinformatics to identify groups of similar biological data points (clusters). Clustering assigns each data point of a given set to one or more of these groups. Here, we focus on the most common case by far, where each data point is assigned to a single cluster; we note that though much of the material in this review extends to the other cases as well.

Clustering has numerous applications in various bioinformatics subfields such as protein function prediction, population genetics, and gene expression analysis. In protein function prediction, clustering can help in grouping proteins based on their structure, sequence, or functional properties. In population genetics, clustering can be used to infer relationships among populations or individuals based on their genetic data. In gene expression analysis, clustering can be used to identify patterns in the expression levels of genes across different conditions, treatments, or patients [1–7]. In this way, clustering can, for instance, help define patient subtypes, i.e., groups of patients with the same expected clinical outcome, which may then allow the prediction of clinical outcomes for new patients. Clustering can also be employed to reduce the large number of genes in gene expression profiles to a smaller number of biologically interpretable gene *modules*, i.e., groups of functionality related genes [1,3,6]. Clustering can provide valuable insights into the relationships and similarities among biological entities and has proven to be a crucial tool in advancing our understanding of biological systems.

In most bioinformatics applications, the input data to clustering is often consistent with multiple, slightly distinct, but equally plausible, clusterings, for several possible reasons: [i] If the input data is considered as a sample from a distribution with a particular global structure, then slightly perturbed input data can also be expected to contain the same global structure, and thus correspond to the same clustering. [ii] Often, multiple models or clustering algorithms, and multiple values of their parameters, are plausible, since, typically, no scientifically precise models of the desired groups, or the relationships between them, are available. [iii] The argument in [i] also holds for multiple input data sets, which can often be considered different samples with the same global structure of interest. In all of the above cases, a set of equally plausible, but often distinct, clusterings stemming from data perturbations, method (parameter) perturbations, or multiple data sets, lead to a set of equally plausible clusterings.

*Consensus clustering* (CC), or *ensemble clustering*, is a class of clustering algorithms that first generates a set of equally plausible results (to which we will also refer as *clusterings*) using a *generation mechanism*. In a second step, CC constructs a clustering whose features are globally as similar as possible to all the features found in the initial (“base”) clusterings, a process referred to as the *consensus mechanism* [8–10]. The underlying idea is that the consensus mechanism is most strongly driven by those features that often occur across base clusterings, and that, therefore, the result can be viewed as robust with respect to the variation in the generation mechanism. Surprisingly, even though the final clustering can be viewed as an average over all base clusterings, it may be radically different from—and, in many cases, more plausible than— any base clustering (see the example in Figure 4 below).

Various possibilities exist for generation and consensus mechanisms. In the following three sections, we review, and structure, the different generation mechanisms in use (Figure 1). As the various consensus mechanisms have already been reviewed elsewhere [8,9], we will focus our discussion on the different generation mechanisms in the specific context of the *co-association matrix* (CM) or *consensus matrix*, as the most commonly used consensus mechanism.

**Figure 1.**
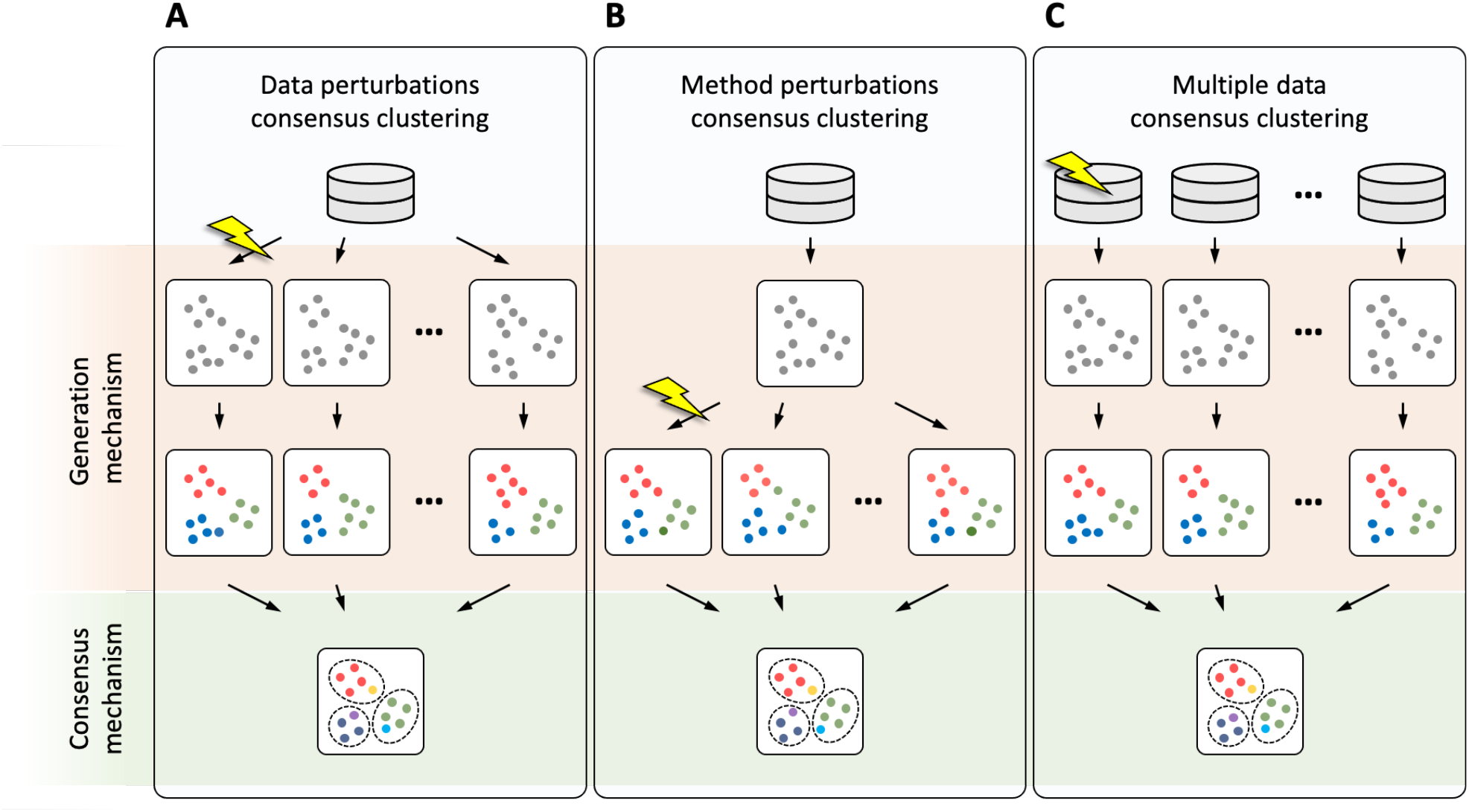
Three types of consensus clustering with distinct generation mechanisms: (A) data perturbation consensus clustering; (B) method perturbation consensus clustering; (C) multiple data consensus clustering

The elements of the CM correspond to the different pairs of data points, each representing the frequency of co-occurrence of both points in the same cluster, across all base clusterings. The CC is then computed using any similarity-based clustering algorithm by treating the CM as a similarity matrix between data points. Algorithm 1 represents CC with CM, as we discuss it in this article.

### Algorithm 1. Generic consensus Clustering Algorithm based on the co-association consensus mechanism

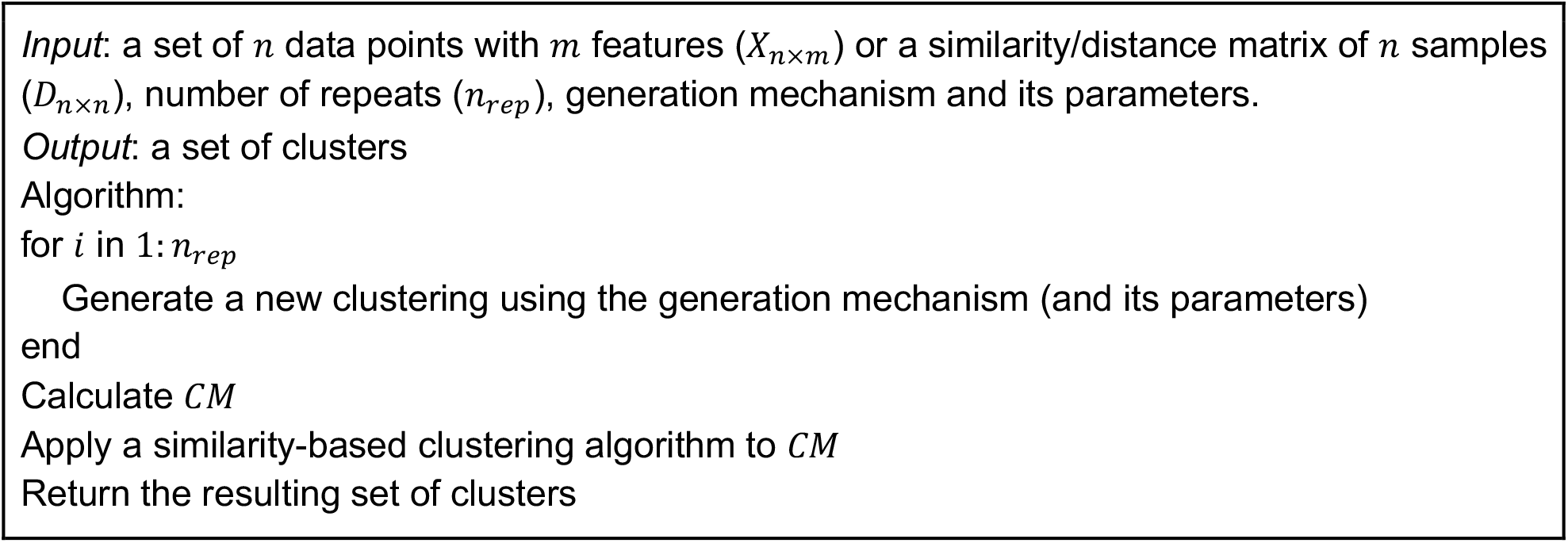

## 2. Data Perturbation Consensus Clustering (DPCC)

To produce a set of distinct, but equally plausible, clusterings, *data perturbation consensus clustering* (DPCC) uses slight perturbations of the input data. The following types of data perturbation have been used in the literature:

- *Addition of noise* [11] perturbs each data point by adding a small offset from a “noise distribution”. The choice of this distribution needs to be adapted to the distribution of the data itself in the different dimensions, and requires assumptions about experimental or empirical noise characteristics.
- *Random subsampling of data features* [12,13] generates new data points through subsampling the set of features. To be used in this context, the subsampling strategy needs to make it likely that enough features are included that characterize the global structure of the data.
- *Random subsampling of data points* [14–17] constrains the input to a subsample of the data. To be effective, subsampling strategy needs to ensure that any pair of data points occurs sufficiently often across the subsamples.

Note that the high computational running time associated with DPCC (and also other forms of CC) can be mitigated through optimizations. For instance, in the case of random subsampling and a distance-based clustering algorithm used in the generation phase, all pairwise distances can be precomputed, instead of recomputing them for each random subsample.

### 2.1 The stability score to estimate the number of clusters

DPCC was originally proposed by Monti et al. [14]. In the same paper, Monti et al. also used the base clustering to formulate a new criterion for the number of clusters, a problem whose formalization is notoriously difficult. Generally, DPCC first uses a base clustering algorithm to produce sets of clusterings with *k* clusters, where *k* varies across a predefined range of values. The final *k* is then chosen as the one with the globally most *stable* clustering, i.e., the clustering that brings together the largest number of pairs of data points that are consistently clustered together across all the data perturbations, which is expressed using a stability score *S*_*k*_ (Algorithm 2).

#### Algorithm 2. The DPCC algorithm by Monti et al. uses the stability score to determine the number of clusters

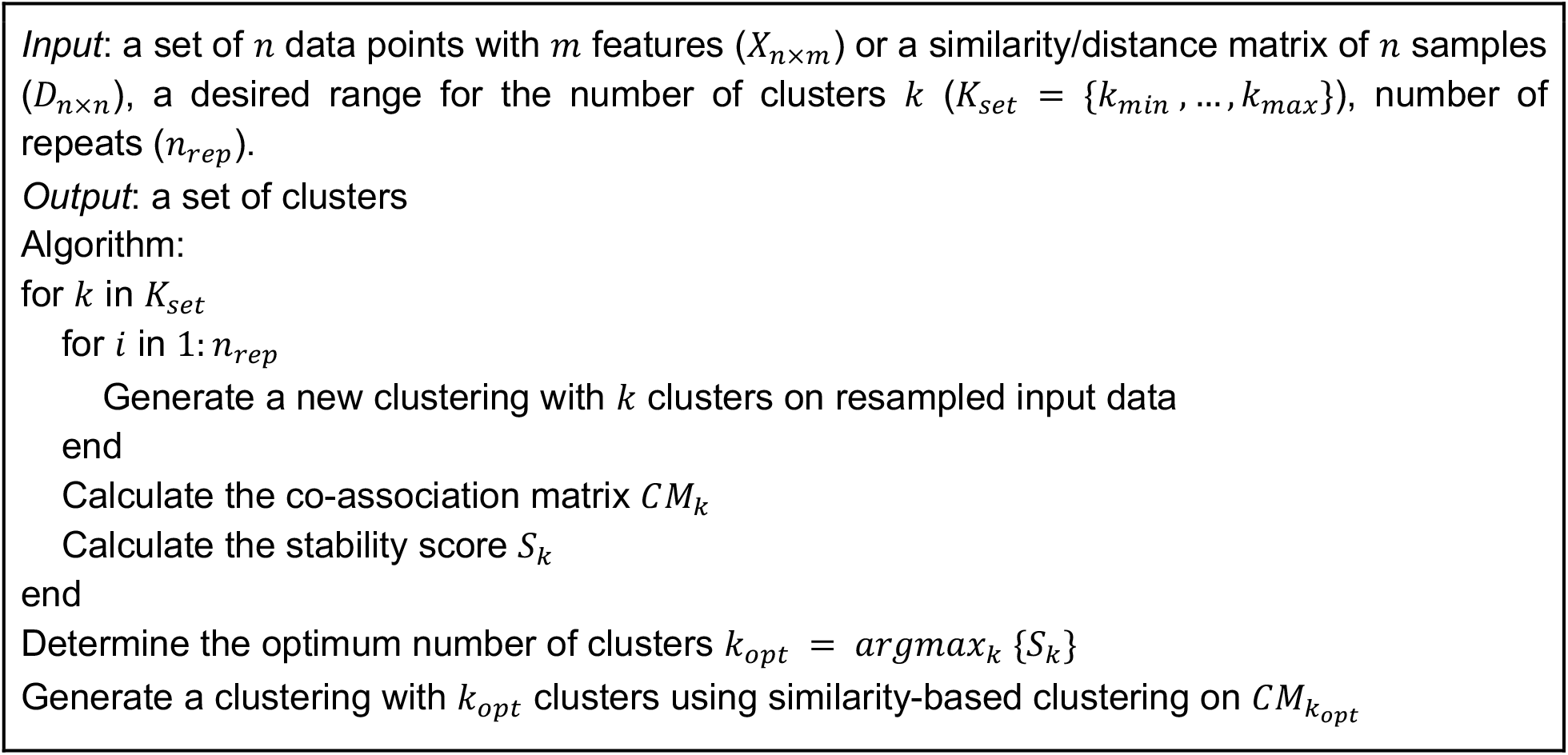

Monti et al. [14] also pointed out the importance of visual analysis of the CM to provide more insight behind the cluster stability. To illustrate their point, we simulate data points of four two-dimensional Gaussian clusters (Figure 2A), and obtain the CM by 100 perturbations, each using random subsampling of 70% of data points followed by *k-medoids clustering* (or *partitioning around medoids* — PAM) for *k*= 2,…, 6. Figures 2B-F show the resulting co-occurrence matrices *CM*_*k*_. Recall that *CM*_*k*_[*i,j*]represents the frequency of data points *i* and *j* being co-clustered. Perfectly consistent base clusterings result in a CM of only zeros and ones, as in the case of *CM*_4_ (Figure 2D). Stability scores therefore capture the tendency of the CM to contain values close to zeros and ones. For this purpose, stability scores often use the *empirical cumulative distribution function* (CDF). Figure 2G shows the CDFs for the matrices *CM*_2_,…, *CM*_6_. Note that the CDF of *CM*_4_ shows sharp jumps around 0 and 1 and a plateau in the middle.

**Figure 2.**
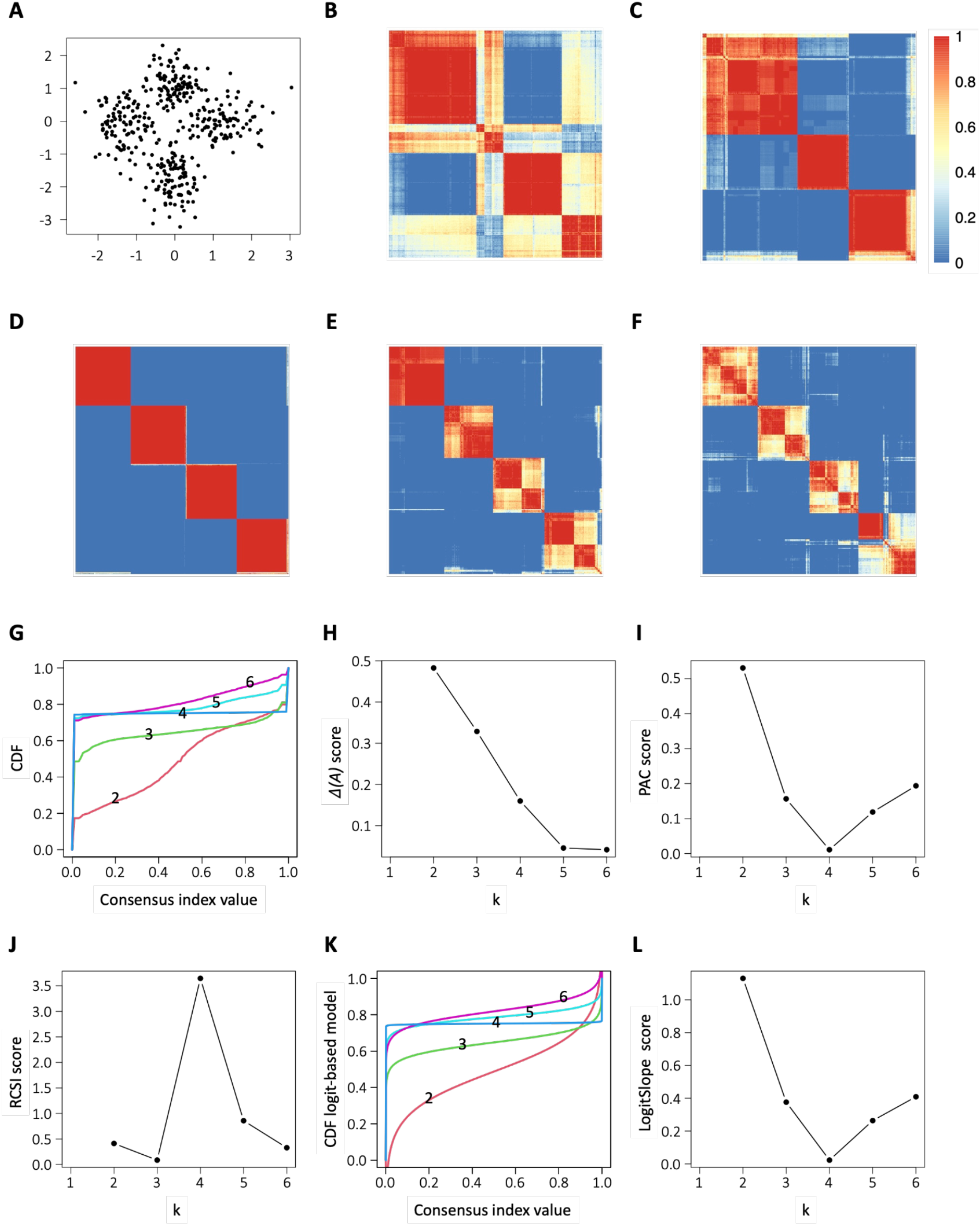
Data perturbation consensus clustering using random subsampling. (A) Input data consisting of samples from four 2D Gaussian distributions. (B-F) Consensus matrices *CM*_2_,…, *CM*_6_. (G) Empirical cumulative distribution function (CDF) of *CM*_2_,…, *CM*_6_. (H-J) stability scores of Δ(*A*), PAC score, and RCSI for *k*∈ {2,…, 6}. (K) Fitted CDFs using *logit* function regression. (L) *LogitSlope k*∈ {2,…, 6}.

Various stability scores have been defined based on the CDF, and applied to estimate the optimal number *k* of clusters. The Δ(*A*) method by Monti et al. [14] uses the relative increase in the area under the CDF between successive value of *k* as the stability score (Figure 2H). Monti et al. also pointed out the possibility to apply stability scores not globally, but at the level of individual clusters. Δ(*A*) has later been shown to have limited accuracy in simulated data [15,16]. Senbabaoglu et al. [15], instead, introduced the *proportion of ambiguous clustering* (PAC) score, which measures the increase of the CDF in its middle segment (Figure 2I). Lower values of PAC are associated with more robustness; thus, the optimum *k* corresponds to the lowest PAC. Senbabaoglu et al. showed that the PAC score improves upon Δ(*A*), but also the alternatives of *CLEST [17], GAP statistic* [18] and *average silhouette width* (ASW) [19]. Further, John et al. [16] proposed a *Monte Carlo reference-based consensus clustering* (M3C) algorithm to improve the PAC score by testing against the null hypothesis of the data consisting of only a single *Gaussian* cluster. Their method is based on a p-value for each *k* that can not only be used to distinguish whether the data has a unimodal Gaussian distribution, but that can also be considered as a stability score (referred to as M3C-Pval). As an alternative stability score, John et al. [16] also proposed the *relative cluster stability index* (RCSI), which corresponds to log-ratio of the mean PAC across randomly generated samples to the PAC of the original data (Figure 2J). They showed that RCSI improves upon the above methods, as well as *Progeny clustering* [20] and *Cophenetic coefficient* [21]. Finally, Lu et al. [22] used the average of the CM values (which we denote by *CMavg* below) as a stability score to compare the stability of different clustering algorithms. The formal definitions of these stability indices are provided in *Supplementary Methods*.

One drawback of the PAC score is that, instead of considering the whole CDF, it only takes a center segment of it into account. We noticed that the CDFs in our case study (Figure 2G) resembles instances of the *logit* function (Figure S1 gives examples), and also included a new stability score based on a *logit* regression model for the CDF of CMs. The *logit* models derived from the empirical CDFs in Figure 2G are shown in Figure 2K. We then simply used the slope in the center of the *logit* function as the *LogitSlope* stability score (Figure 2L – for details, see *Supplementary Methods*).

### 2.2 Empirical comparison of stability scores

To assess the ability of different stability scores, including Δ(*A*), PAC, LogitSlope, M3C, and ASW (based on *k-means* clustering) to estimate the number of clusters, we generated synthetic data from a model with different numbers of well-separated clusters (see *Supplementary Methods* for details). Briefly, given a number *k*_*sim*_, our simulation model samples *k*_*sim*_ “ground truth” clusters around centers *i* = 1,…, *k*_*sim*_ at a distance of *r* of the origin, and at an angle of 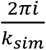, where *r* is drawn from a uniform distribution over the real interval [1,2](Figure S2). The points of each cluster are then sampled from a two-dimensional Gaussian distribution whose mean is the cluster center. The diagonal entries of the symmetric covariance matrix are drawn from a uniform distribution over 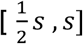, and all other entries are drawn from a uniform distribution 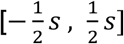. This choice ensures that the covariance matrix is positive definite, and thus the ground truth cluster is not degenerate (Figure S3). The parameter *s* controls the overall level of overlap between ground truth clusters (*cf. Supplementary Methods*).

In our analysis, we performed DPCC with the PAM as the base clustering method, and estimated the estimated number *k*^*∗*^ of clusters to test different stability scores. Figure 3 and Figure S4 show the accuracy of the estimate *k*^*∗*^ of *k*_*sim*_ by different stability scores, and under different values of *s*. As expected, the accuracy generally tends to decrease as *s* increases. Overall, we observe that LogitSlope provides the highest accuracy in almost all cases, closely followed by PAC and M3C-RCSI. The other methods perform consistently worse.

**Figure 3.**
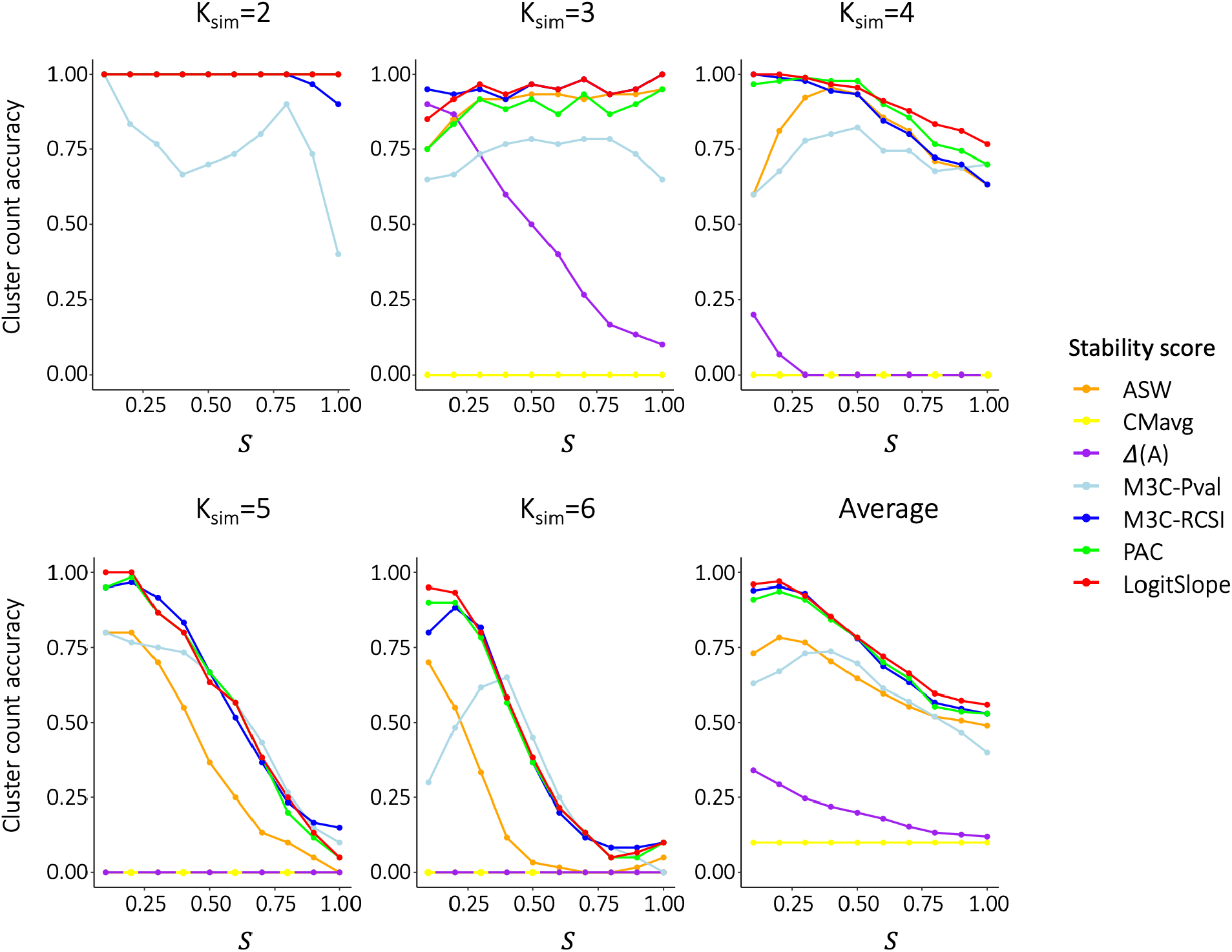
Empirical accuracy of different stability scores to estimate the “ground truth” number *k*_*sim*_ of clusters on simulated single Gaussian clusters. Note that, in the case *k*_*sim*_ = 2, all scores (except for M3C-RCSI and M3C-Pval) achieve a constant accuracy of 1.

**Figure 4.**
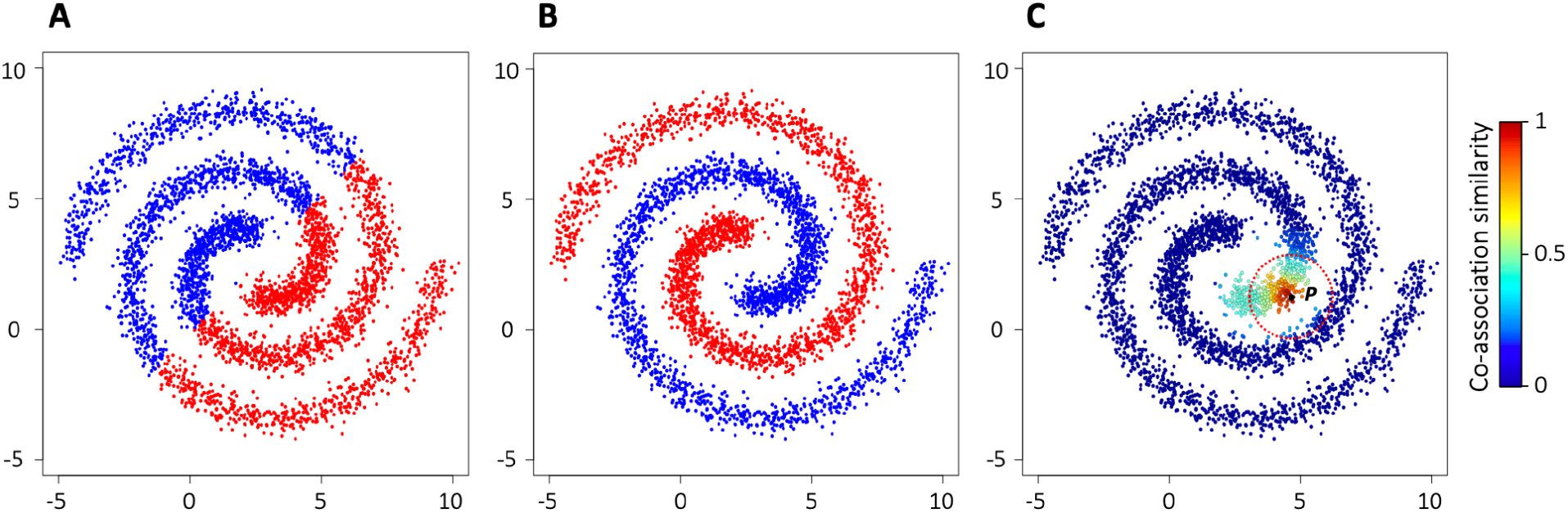
(A) *k*-means clusters using *k*= 2. (B) Consensus clustering using a range of values for *k*∈ [10,100]. (C) Consensus similarity between a point *P* and all the other points

## 3. Method Perturbation Consensus Clustering (MPCC)

*Method perturbation consensus clustering* (MPCC) is based on the idea of generating different equally plausible base clusterings by varying the clustering algorithm [23–26], hyperparameters [27–31], algorithm initializations [27,32], preprocessing methods [33,34] or combinations of these aspects [9,28,35].

### 3.1 Stability against hyperparameter variation

“Hyperparameter” typically designates method parameters that do not depend on input data, but global parameters that are set, independently of the input data, by the user. Examples of hyperparameters include noise parameters, the expected shape of clusters, parameters of the architecture of a neural network model, or prior distributions in Bayesian analysis. In those cases where large amounts of data are available, *hyperparameter tuning* can be used to select parameters in a data-driven way (such as the number *k* of clusters in Section 2.1). But, often, neither data nor knowledge of the application suggests a particular value of *k* that would be clearly better than other values. Also in these cases, CC with base clusters from a range of hyperparameter values may be an effective means to obtain robust results.

Even though CC aims to produce a clustering that is globally similar to the base clusterings at the level of features, the result can be drastically different at the level of the clustering. We illustrate this point using an MPCC that generates base clusterings with a varying number of clusters [8,27,28,35,36]. In the generation phase, we use *k*-means for the base clustering algorithm, and at each iteration, we randomly choose *k* from a predefined range, and lastly calculate the CM (Algorithm 3).

#### Algorithm 3. Calculating co-association matrix based on k-means

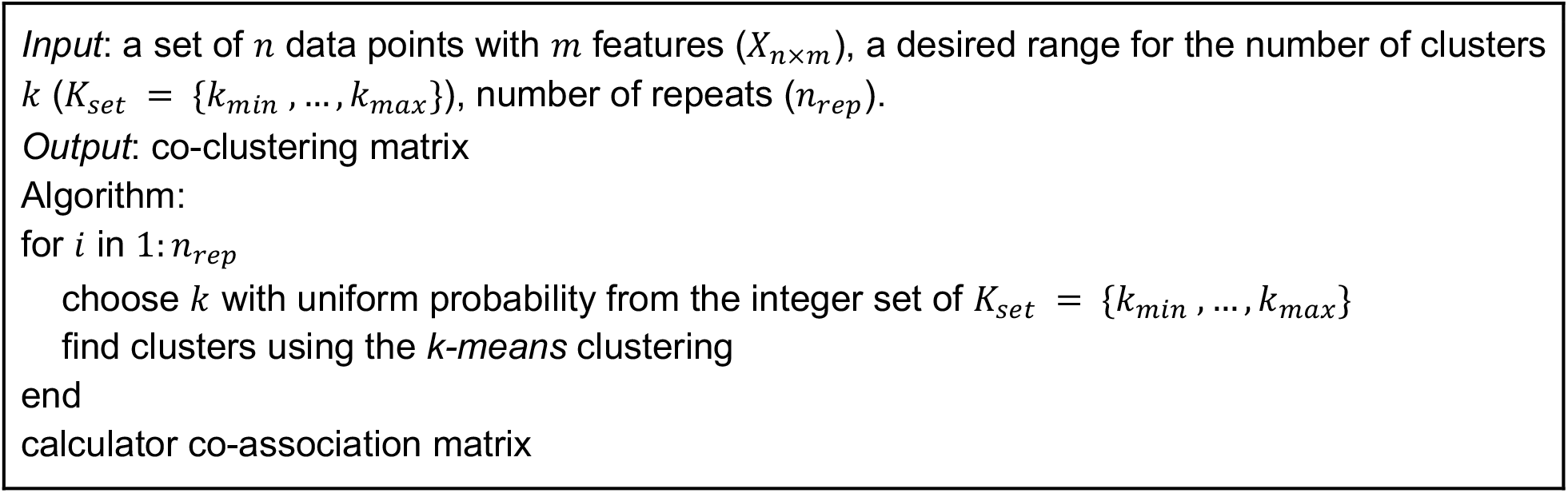

Figure 4A shows the result of CM-based CC on the “Spiral” dataset, based on *k*-means clustering with *k*= 2. Figure 4B shows the result of MPCC, using *k*-means base clusterings over a range of *k*, followed by hierarchical clustering on the CM. Interestingly, even though all base clusters occupy convex regions, the regions occupied by the two clusters of the resulting CC are highly non-convex. Figure 4C provides a statistical argument for why this happens. First, note that most points within a fixed small Euclidean distance from an arbitrary, fixed, data point *P* (dashed circle) are on the same spiral arm. Thus, it is these points on the same spiral arm that tend to co-occur together in the base clusterings, and thus contribute to a high co-association similarity between them. Figure 4C shows the values of the co-association similarities between *P* and the other data points using colors.

In the above example, co-association similarity is thus able to distinguish pairs of points on the same spiral arm from other pairs that have the same Euclidean distance from each other, but do not lie on the same spiral arm (Figure 4C). Co-association similarity thus functions as an alternative to Euclidean distance as “locally structure-aware” similarity measure. We believe that co-association similarity can also be used in many other cases of potentially more complex data geometries, and beyond the context of clustering.

### 3.2 Stability against initializations

Many clustering methods, such as those based on matrix factorization [37,38], artificial neural networks [39,40], and model-based clustering [41], require iterative methods to solve non-convex optimization. Starting from distinct, often random, initial conditions, these methods often converge to a distinct local minima. To make these methods stable against the randomness of the initial conditions, the clusterings generated from different initial conditions can be used as input for CC [32].

### 3.3 Stability against pre-processing methods

Clustering results in bioinformatics commonly also depend on specific methods selected for data preprocessing. For many types of data, such as DNA sequence, transcriptome, proteome, or metagenome data, no obvious single best preprocessing method exists. As one example, for the case of gene expression data, typically, multiple normalization approaches have been judged to be equally plausible [33,34,42]. Different processing methods then often lead to multiple equally plausible inputs for consensus clustering.

## 4. Multiple Data Consensus Clustering (MDCC)

A third type of CC is *multiple data consensus clustering* (MDCC), which is based at base clusterings generated from multiple datasets. Clusters can consist of either samples or features (*e*.*g*., genes), corresponding to the two distinct scenarios [43], as shown in Figure 5: [A] *multi-view CC*, where samples are clustered according to their similarity across distinct sets of features (e.g., transcriptomics and proteomics, other instances of multi-omics analysis), and [B] *multi-cohort CC*, where features (e.g., genes) are clustered according to their similarity across different sets of samples^1^.

**Figure 5.**
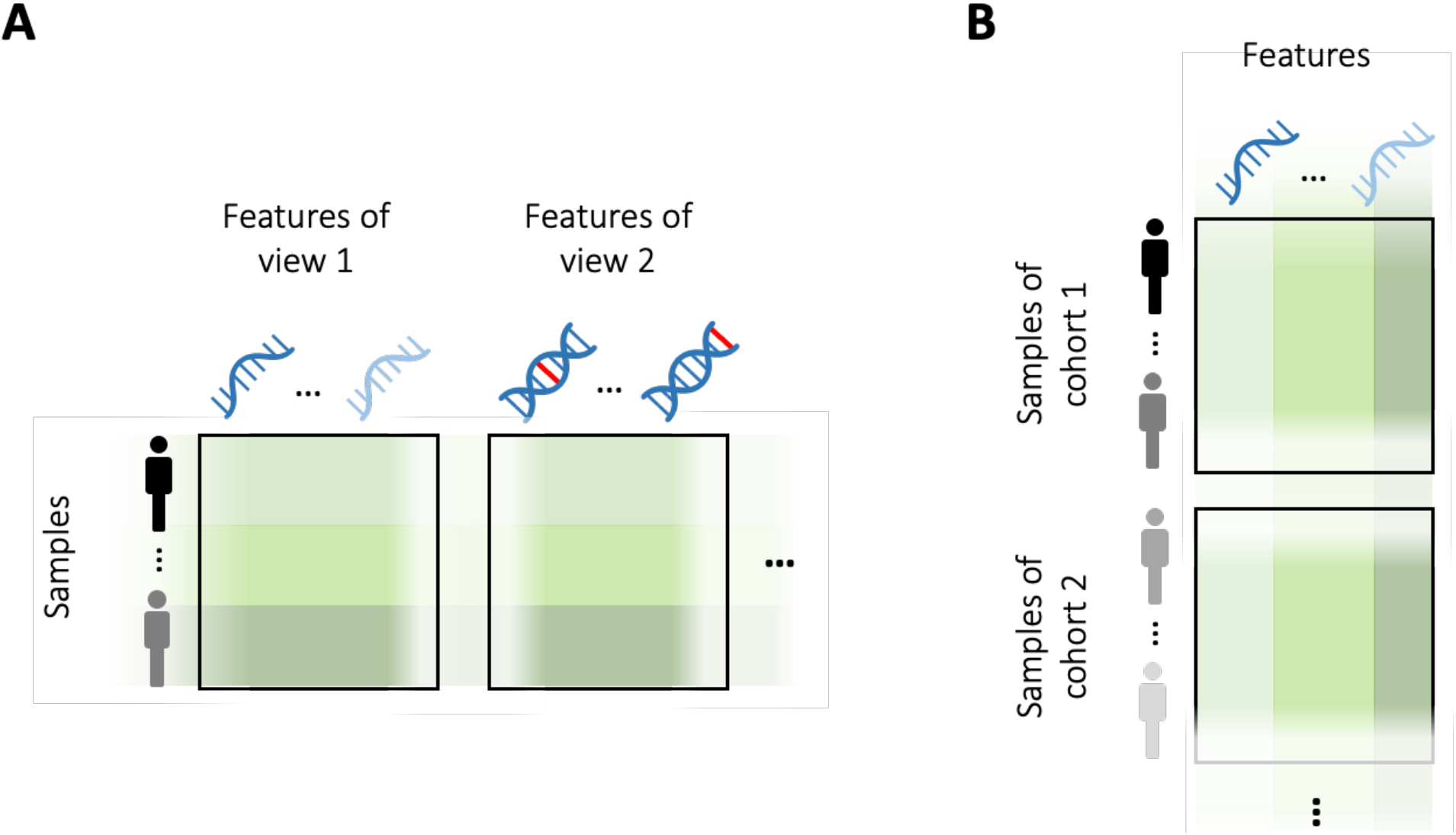
Two distinct scenarios for multi-data consensus clustering: A) multi-view consensus clustering and B) multi-cohort consensus clustering

Because of the presence of, often, uncharacterized specific biases of individual datasets, multi-cohort CC may result in more robust biological results than the analysis of individual cohorts. In the following case study, we compare *gene modules* (i.e., clusters of genes) derived from individual datasets to the gene modules obtained from multi-cohort CC. We used the publicly available transcriptomic data from five *systemic lupus erythematosus* (SLE) cohorts in the *gene expression omnibus* (GEO) database. To derive gene modules, we applied PAM clustering to each dataset separately, and then applied multi-cohort CC to the same five datasets (details described in *Supplementary Methods*). The number of clusters was set to four based on the LogitSlope stability score (cf. Section 2). To interpret the resulting gene modules in the context of SLE, we used over-representation analysis on the gene expression modules from Chaussabel et al. [45] and Li et al. [46] using the R package *tmod* [47].

Figure 6 shows all enriched gene expression modules across all clusterings. Note that the CC result is enriched much more clearly in pathways corresponding to distinct established immunological facets of SLE, and relevant cell types, than the cohort-specific clusterings. The set of interferon gene modules are detected with almost perfect sensitivity and specificity by MDCC, but not by any of the cohort-specific clusterings. Also, note that certain gene modules in the “Lymphocyte” category are *only* enriched in CC. These observations suggest that MDCC is superior clustering of on individual datasets for pinpointing the true molecular basis of SLE.

**Figure 6.**
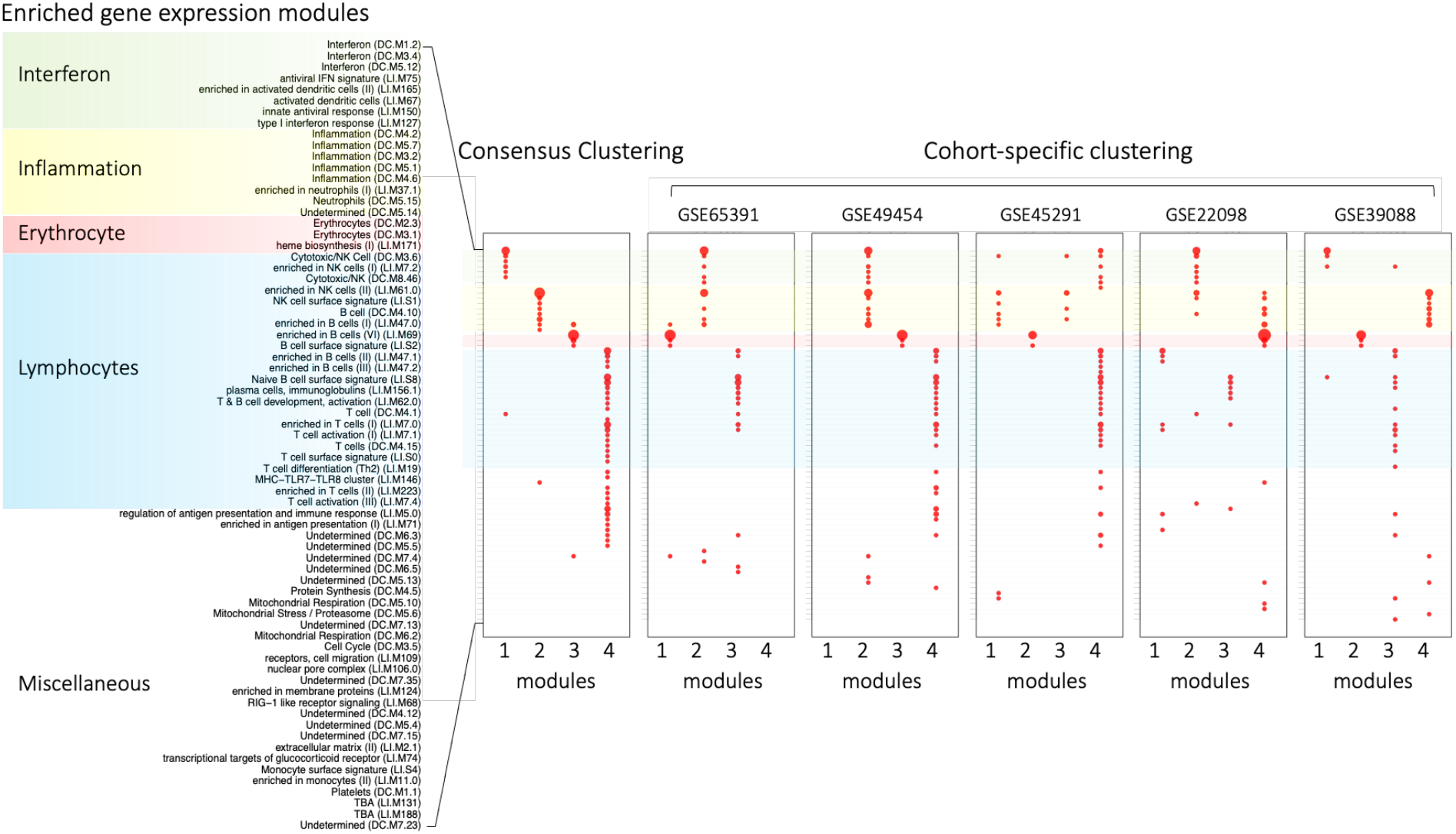
Enriched gene modules found in cohort-specific, and consensus clusters of blood transcriptomes of five different SLE cohorts. Multi-cohort CC much more consistently identifies biological processes and cell types known to be related to SLE. Dot size is proportional to negative enrichment log *p-value*, smallest dots corresponding to *p*=0.05.

Note that MDCC can be implemented in a distributed manner that is highly compatible with data privacy concerns and restrictions [35,44]. The acquisition and storage of biomedical (and especially clinical) cohort data are not only time-consuming and costly but diverse datasets also can often not be merged due to privacy and ethical concerns, even when anonymized [48]. Since most stages of MDCC can be performed using distributed computing, only information about the cluster membership of the data points needs to be merged in the final consensus phase of MDCC (cf. Figure 1). MDCC is therefore also applicable in situations where data cannot be shared due to competition between the owners of different datasets.

## 5. Discussion

Even though most classical clustering algorithms provide a single clustering as a result, in most bioinformatics applications, other, often equally likely, clusterings exist. In this article, we discussed three types of variation that can lead to multiple, distinct, candidate clusterings: [i] the input data itself that comprises technical variation in the input data, [ii] the clustering method and its parameters, which are often chosen in an *ad hoc* fashion, and [iii] biological idiosyncrasies of particular datasets, which are often uncharacterized. While the candidate clusterings are usually somewhat distinct, they often share certain *consensus features*, such as the co-occurrence of certain input points in a cluster. These features can then be used to generate a consensus clustering (CC) that is, globally, maximally similar to the set of candidate clusterings. CC can be interpreted as a principled extension to robustify any basic clustering algorithm.

In this article, we presented the first systematic overview of CC algorithms tailored to the above three sources of variation, with a focus on co-occurrence as the most commonly used consensus feature. Correspondingly, we distinguish three classes of CC methods: data perturbation CC (DPCC), method perturbation CC (MPCC) and multiple data CC (MDCC). DPCC clusters are robust against data perturbations caused by measurement error and/or sampling method, while MPCC provides robustness against particularities of the computational pipeline, which can include choice of the method, its hyperparameters, and its probable stochasticity. MDCC yields clusters that are robust against idiosyncrasies of individual datasets. Together with available software (including the approaches discussed in this review, see Code Availability below), CC provides a universally and easily applicable functional extension of basic clustering algorithms that can be used to increase the quality of bioinformatic data analysis.

We mention a few relevant extensions to the CC methodology and open research questions. Firstly, prior knowledge can be incorporated into the consensus phase by augmenting the data-driven set of base clusterings with knowledge-driven clusterings of networks [35,49]. To mention one such possible augmentation in a bioinformatic application, one can use a network of gene pairs that are known to co-occur in specific biological pathways, or known to functionally interact. Such a network, e.g., the STRING database [50], can then be added as an additional “knowledge-based” component to the CM. We note in passing that, on a technical level, the use of networks (as opposed to the more constrained use of clusters) in the consensus phase of CC is an extension of the way we introduced CC in Section 1, but that this extension is technically straightforward, and may be well justified for other purposes as well. The different types of CC can also be combined to confer robustness to multiple types of variation. In its simplest form, all three generation mechanisms could be used to generate a large set of base clusterings, or different types of CC could be applied serially.

In bioinformatics applications, other types of robustness than the ones we discussed so far are important, including statistical robustness to outliers and the violation of distributional assumptions. We are not aware of the use on these types of robustness for CC, and believe that the sensitivity of certain clustering algorithms to outliers and the violation of distributional assumption would warrant their inclusion. To incorporate robustness against this, and other, types of variation into CC, there are two (non-exclusive) options. On the one hand, one can use, in the generation phase, base clustering algorithms that are robust against these types of variation. On the other hand, custom types of data perturbation could be employed in the CC generation phase, e.g., by adding outliers from a hypothetical outlier distribution or by resampling the input data according to non-canonical distributions.

For any type of generation mechanism, it is important to control the level of diversity of the generated clusterings. If the base clusterings are not diverse enough (i.e., they all produce similar results), the consensus clustering may not offer any improvement. Conversely, if the base clusterings are too diverse or inconsistent, the consensus result may not make sense or may obscure the true structure in the data. How to control the level of diversity of the generated clusterings is an important question in practice that we did not see addressed so far.

Finally, while stability scores, as introduced, can be used to assess the stability and robustness of clusters, statistical testing can provide important complementary information about the likelihood of the observed clusters being meaningful and non-random, and can also been used to determine the optimum number of clusters [16,51]. The choice of an appropriate null model is important to obtain a good approximation of the biological validity of a clustering (or individual clusters), but often not obvious.

## Code Availability

We provide open-source *R* codes that allow the reader to recapitulate the examples in this review, and to apply the different types of consensus clustering to other data. These codes include functions for consensus clustering (DPCC, MPCC and MDCC), stability score calculation (LogitSlope, PAC, Δ(*A*), CMavg), and the generation of simulated data (single- and mixture-Gaussians) and are available via the GitHub repository:

https://github.com/behnam-yousefi/ConsensusClustering

## Footnotes

1 These two scenarios have been called *horizontal clustering* and *vertical clustering [43]*. However, we prefer *multi-view*/*multi-cohort clustering*, since the interpretation of these terms depends on the, often arbitrary, choice of data representation in any specific case has led to confusion in several cases [36,44].

