## Supplementary Methods for "Consensus Clustering for Robust Bioinformatics Analysis"

#### 1. Synthetic “Ground truth cluster” model for the empirical comparison of stability scores

In our simulation model, we generate $k_{sim}$ clusters of $n_{mixture}$ Gaussians each. $n_{mixture}$ is either equal to 1 (“single Gaussian clusters”), or drawn from the set $\{2,3,4\}$ (“multiple Gaussian clusters”) with equal probabilities. To generate the clusters, we first determine the cluster centers

$$\boldsymbol{c}_i = \left[c_{x,i}\,, c_{y,i}\right]^T \ ; \quad i = 1, \dots, k_{sim}$$

where

$$c_{x,i} = r_i \cdot \cos(\frac{2\pi i}{k}),$$

$$c_{y,i} = r_i \cdot \sin(\frac{2\pi i}{k}), \text{ and}$$

$$r_i \sim uniform(r_{min}, r_{max}).$$

This is illustrated in Figure S2A and S2B. For our case studies in Section 2, we chose $r_{min} = 1$ and $r_{max} = 2$. In case of $n_{mixture} = 1$ (single Gaussian cluster), we generate random Gaussian points around each center with the covariance matrix

$$\Sigma_i = \begin{bmatrix} a_{i,11} & a_{i,12} \\ a_{i,21} & a_{i,22} \end{bmatrix}$$

where

$$a_{i,11}\,, a_{i,22} \sim uniform(\frac{s}{2}, s)$$

$$a_{i,12} = a_{i,21} \sim uniform(-\frac{s}{2}, +\frac{s}{2})$$

The elements of the covariance matrix are designed in a manner that guarantees a positive definite matrix. Note that the parameter $s$ controls the overall level of overlap between ground truth clusters.

Moreover, each cluster $i$ can be composed of a mixture of $n_{mixture}$ Gaussians, each centered around a sub-center, whose position is slightly modified by adding an offset to $\boldsymbol{c}_i = \left[c_{x,i}, c_{y,i}\right]^T$ resulting in $n_{mixture}$ points as $\boldsymbol{c}_{ij} = \left[c_{x,i,j}, c_{y,i,j}\right]^T$ $i = 1, \ldots, k_{sim}$ and $j = 1, \ldots, n_{mixture}$, where

$$c_{x,i,j} = c_{x,i} + \varepsilon_{x,i,j}$$

$$c_{y,i,j} = c_{y,i} + \varepsilon_{y,i,j}$$

$\varepsilon_{.,i,j} \sim uniform(-sd_{max}, sd_{max})$,

as shown in Figure S2A, S2C.

In our comparison of different stability scores, we increased the parameter $s$, that controls the standard deviation of each cluster point from its center in each dimension, from 0.1 to 1 in increments of 0.1. The lower limit of 0.1 was chosen to ensure a sufficient degree of separation between the clusters: note that the minimum possible distance $d_{min}$ between any two cluster centers is

$$d_{min} = 2 \cdot r_{min} \cdot sin\,(\frac{\pi}{k})$$

In our study, $k = 6$ leads to the minimal value of $d_{min} = 1$. We chose the minimum value for the parameter $s$ as $\frac{d_{min}}{10}$. Figure S3 illustrates the density between a pair of closest clusters. As, even in this extreme case, the density between the clusters drops significantly, we believe that all $k$ clusters can be considered to be well-separated, and hence our simulation data can be considered to consist of $k$ clusters.

#### 2. Co-association matrix

The co-association matrix (CM) is the key component of the most common consensus mechanism used for consensus clustering. In the first step, different base clusterings are obtained through the generation mechanism by data perturbations, method perturbations, and multiple datasets. Next, $CM[i, j]$ is defined as the frequency with which samples $i$ and $j$ are co-clustered across the base clusterings. Note that a CM corresponding to perfectly identical base clusterings contains only zeros and ones; the stability score quantifies how distinct the base clusterings are from this. Monti et al. (2003) chose to define a stability score using the *empirical cumulative distribution* (CDF) of CM elements over the range $[0, 1]$ as

$$CDF(x) = \frac{1}{n(n-1)/2} \sum_{i<j} u(M[i, j] - x)$$

where $u(x)$ is the *Heaviside* function (or step function) defined as

$$u(x) = \begin{cases} 1 & if\ x > 0 \\ 0 & otherwise \end{cases}$$

and $n$ is the number of samples.

#### 3. Stability scores

The CM can be used to calculate a stability score, which can then be used to choose specific methods or hyperparameters, most commonly, the number of clusters. In the following we describe the stability scores we used in this article.

***Δ(A).*** Monti et al. (2003) defined the stability function *Δ(A)* that is increasing in the number *k* of clusters, and proposed that the relative increase in the area under the CDF (denoted as *A*), from $k$ to $k+1$ can be used to determine the most suitable $k$. The relative increase in Δ(*A*) is calculated as

$$\Delta(A)_k = \begin{cases} A_k & if\ k = 2 \\ \dfrac{A_{k+1} - A_k}{A_k} & if\ k > 2 \end{cases}$$

and the optimum $k$ is one where $\Delta(A)_k$ is maximal.

*PAC score*. Senbabaoglu et al. (2014) defined the *proportion of ambiguous clustering* (PAC) score as the fraction of CM values falling in the intermediate sub-interval $(x_1, x_2)$ where $x_1, x_2 \in [0, 1]$ (by default $x_1 = 0.1$, and $x_2 = 0.9$).

$$PAC(x_1, x_2) = CDF(x_2) - CDF(x_1)$$

The PAC score thus represents the increase of the CDF in its middle segment. Lower values of PAC are interpreted as a higher degree of robustness; thus, the optimum $k$ corresponds to the lowest PAC. One limitation of the PAC score is that, instead of considering the whole shape of the CDF, it only takes its middle segment into account.

*LogitSlope.* The *LogitSlope* stability score, which we introduce in this article, models the distribution of values in the CM as a $logit(.)$ function,

$$logit(x) = \log\left(\frac{x}{1-x}\right)$$

$$CDF(x) \approx a \cdot Logit(x) + b$$

We determine $a$ and $b$ by fitting the empirical CDF of the CM using linear regression and define $a$ to be the *LogitSlope* stability score. Note that the slope of our *logit* model in the center of the domain [0,1] is 4*a*. Figure S1 illustrates how lower values of *a* correspond to sharper jumps around 0 and 1 and flat plateaus in the middle segment, and thus, to more stable clusterings.

*Reference-based CC.* John et al. (2020) proposed the M3C method to improve upon the PAC score. They used a Monte Carlo simulation-based approach to generate samples from a single Gaussian cluster $n$ times, and calculated the empirical null PAC score distribution ($PAC_{null}$). Next, they calculated the empirical PAC score for each $k$ ($PAC_k$). Accordingly, the $RCSI_k$ is calculated as

$$RCSI_k = \log_{10}\left(\frac{1}{n}\sum_{i=1}^{n} PAC_{null\ i}\right) - \log_{10}(PAC_k)$$

John et al. (2020) further calculated a Monte Carlo p-value for each $k$, under the null hypothesis that the PAC score originates from a single Gaussian cluster, as

$$P_k = \frac{o_k + 1}{n + 1}$$

where $o_k$ is the number of times $PAC_{null}$ is less than or equal to $PAC_k$. M3C-based stability indices are calculated using the code available in https://github.com/crj32/M3C (John et al. 2020).

#### 4. Gene module detection in SLE

We downloaded blood transcriptome profile of the following five different datasets of SLE that are publicly available through gene expression omnibus (https://www.ncbi.nlm.nih.gov/geo): GSE65391, GSE49454, GSE45291, GSE22098, GSE39088. Background correction and the between-array normalization were performed using the *limma* R package (Ritchie and Phipson, 2015) on the probe level, and probes were annotated with their corresponding gene symbol. Among those probes that refer to the same gene symbol, we chose the data from the probe with the highest standard deviations be. For each dataset, any gene not among the 5,000 genes with the highest standard deviation across patients were discarded, and, among those genes that remained, only those common to at least three datasets were selected. Next, the cohort-specific gene co-expression networks (GCNs) were constructed using signed Pearson correlation (Langfelder and Horvath, 2008) To find gene modules, we applied the PAM algorithm to each dataset separately followed by a PAM clustering on the obtained CM.

### Supplementary Figures

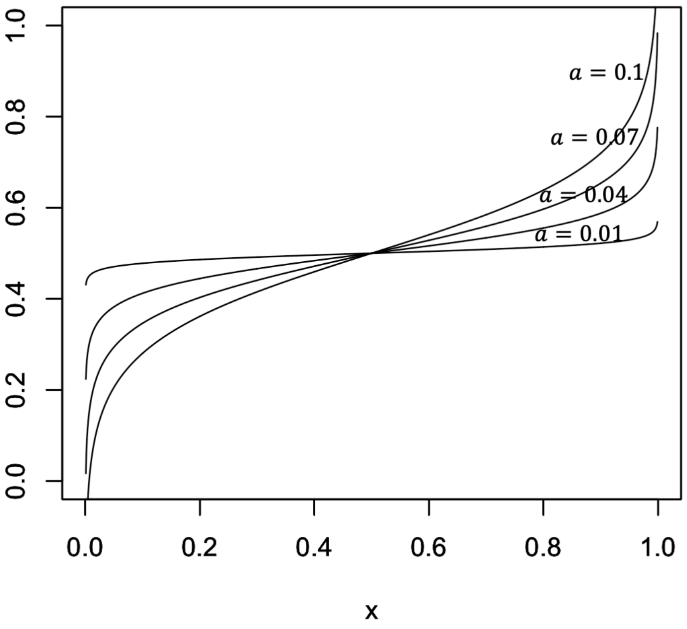


**Figure S1.** Logit functions $a \cdot Logit(x) + 0.5$ used as models for the entries of the co-association matrix CM.

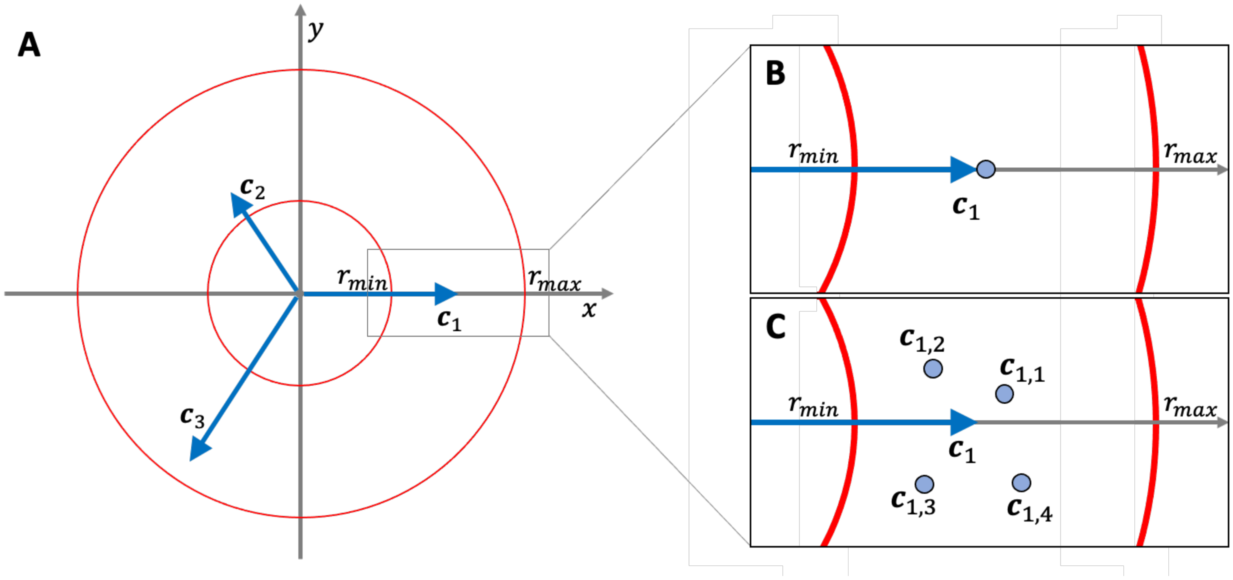


**Figure S2.** Defining the centers of our synthetic cluster generation model for $k = 3$ clusters. (A) Our model places cluster centers $\boldsymbol{c}_i$, $i \in \{1,2,3\}$ at a distance of an angle $\frac{2\pi i}{3}$ and a distance drawn from a uniform distribution over $[r_{min}, r_{max}]$. (B) In case of the single Gaussian clusters, the actual cluster centers are considered as $\boldsymbol{c}_i$ . (C) while in case of the mixture of Gaussian clusters, each $\boldsymbol{c}_i$ is slightly modified by adding an offset to generate $n_{mixture}$ (here $= 4$) new centers $\boldsymbol{c}_{ij}$, $i \in \{1,2,3\}$ and $j \in \{1,2,3,4\}$

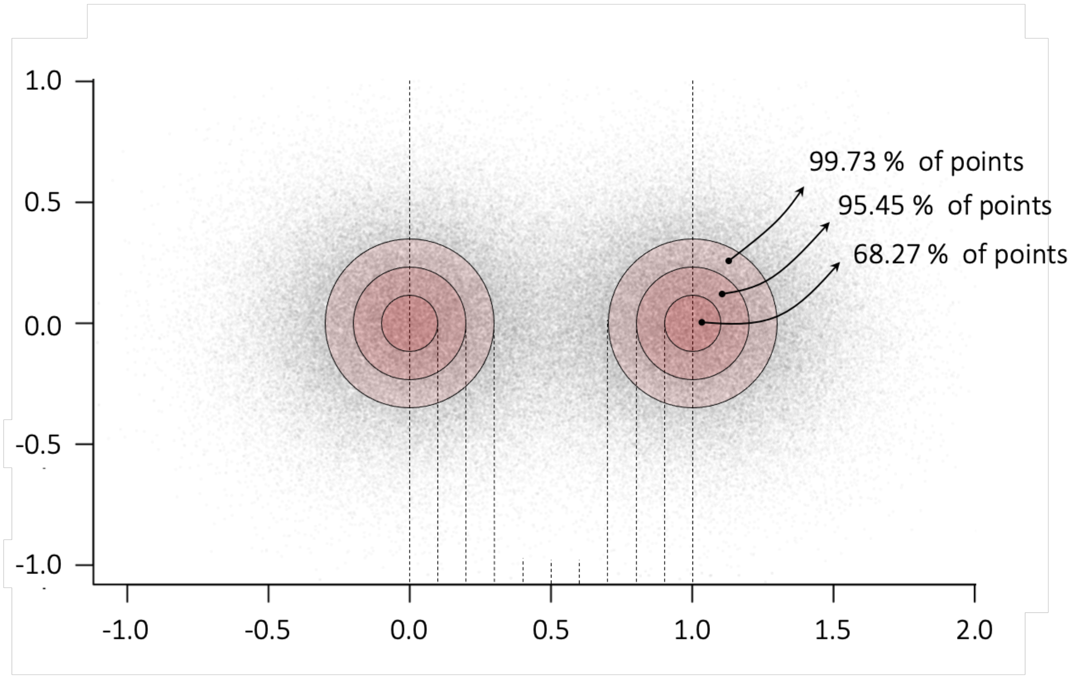


**Figure S3.** The distribution of two clusters from a 2D Gaussian distribution, where the standard deviation $\sigma$ for both coordinates is 0.1, and the clusters are centered at a unit distance from each other. The concentric circles around both cluster centers correspond to distances of $\sigma$, 2 $\sigma$, and 3 $\sigma$ to the respective cluster center.

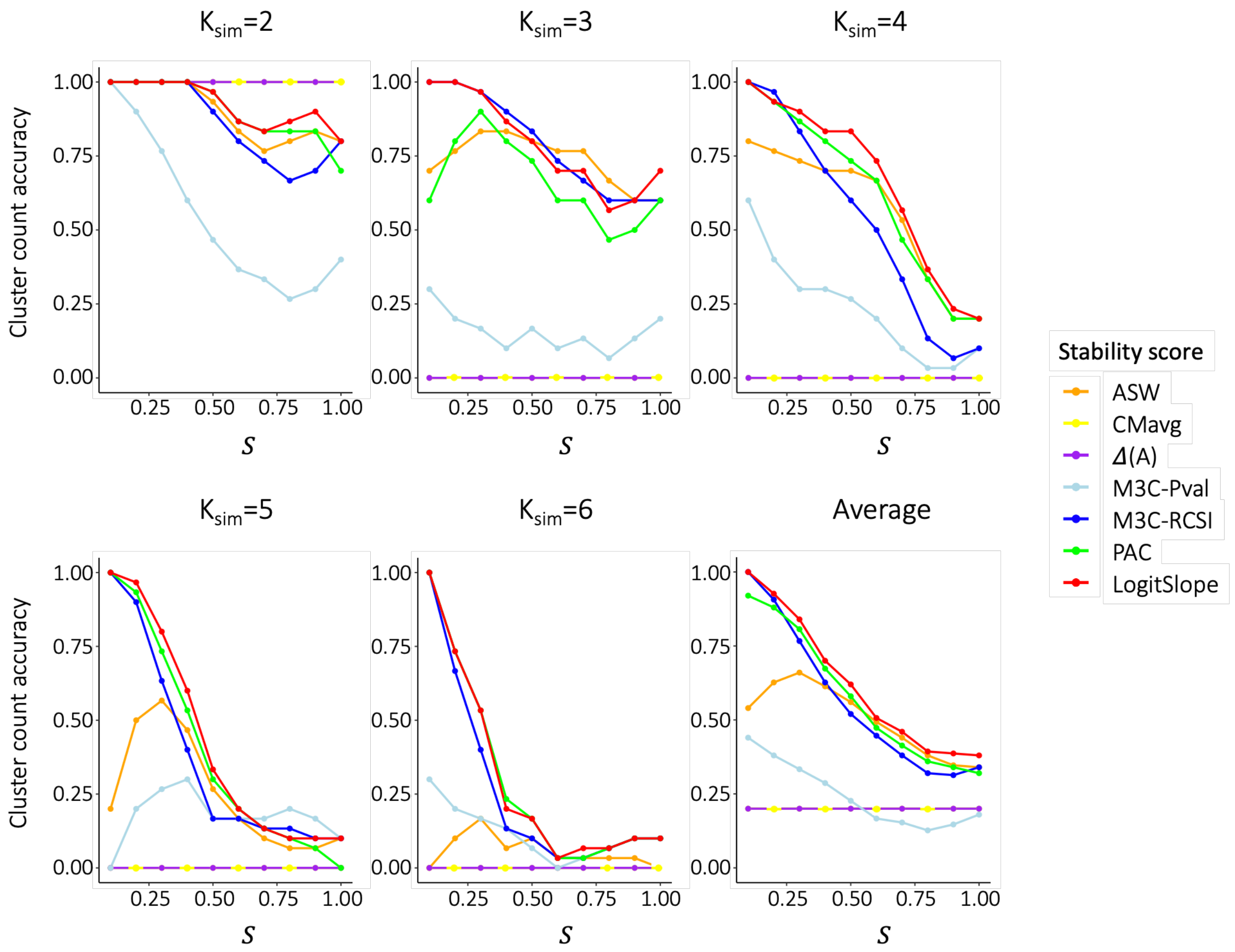


**Figure S4.** Comparison of different stability scores on simulated mixture of Gaussian clusters.
